# Evolutionary Genomics of Sister Species Differing in Effective Population Sizes and Recombination Rates

**DOI:** 10.1101/2023.05.13.540670

**Authors:** Zhiqiang Ye, Michael E. Pfrender, Michael Lynch

## Abstract

Studies of closely related species with known ecological differences provide exceptional opportunities for understanding the genetic mechanisms of evolution. Here, we compared population-genomics data between *D. pulex* and *D. pulicaria*, two reproductively compatible sister species experiencing ecological speciation, the first largely confined to intermittent ponds and the second to permanent lakes in the same geographic region. *D. pulicaria* has lower genome-wide nucleotide diversity, a smaller effective population size, higher incidence of private alleles, and substantially more linkage-disequilibrium than *D. pulex*. Functional enrichment analysis revealed that positively selected genes in *D. pulicaria* are enriched in potentially aging-related categories such as cellular homeostasis, which may explain the extended lifespan in *D. pulicaria*. We also found that opsin-related genes, which may mediate photoperiodic responses, are under different selection pressures in these two species. Additionally, genes involved in mitochondrial functions, ribosomes, and responses to environmental stimuli are found to be under positive selection in both species. Our results provide insights into the physiological traits that differ within this regionally sympatric sister-species pair that occupies unique microhabitats.

## Introduction

Comparative population-genomic analyses of related species with known population-genetic and/or ecological differences provide unique opportunities for obtaining insights into the genetic mechanisms of evolution. Animal species subject to such work include fish (Piechel and Marques 2017; Crawford et al. 2020), mice (Bedford and Hoekstra 2015; Wang and Payseur 2017), and dipterans (Bergland et al. 2016; Bradshaw et al. 2018; Rodrigues et al. 2021). Of special interest is the North American species pair of microcrustaceans, *Daphnia pulex* and *Daphnia pulicaria,* each of which is the other’s closest relative. The two species are nearly identical morphologically (Brandlova et al. 1972), but whereas *D. pulex* mainly inhabits ephemeral ponds, *D. pulicaria* are generally restricted to permanent lakes. Often the bodies of water involved are separated by only a few meters. Given the lack of compelling morphological distinctions, species ascertainment was historically based on allozyme markers, with *D. pulex* isolates typically being homozygous for the slow allele (SS), and *D. pulicaria* homozygous for the fast allele (FF) at the lactate dehydrogenase (*Ldh*) locus (Hebert et al. 1989; Crease et al. 2011). More recently, *>* 20,000 species-specific single-nucleotide markers have been found to distinguish the two species across the genome (Ye et al. 2022).

The cyclically parthenogenetic lifestyle of *Daphnia* results in a strong correlation between habitat specialization and the degree of recombination in the species pair. Throughout the bulk of the growing season, *Daphnia* populations reproduce by ameiotic parthenogenesis, but can switch to sexual reproduction and resting-egg production following the environmental induction of male production. (Although obligately asexual forms of the two species exist, here we focus entirely on populations capable of sexual reproduction). As occupants of temporary ponds, the *D. pulex* populations referred to in this study typically reproduce asexually every 3 to 5 generations, and consequently experience as much if not more recombination per generation than obligately sexual species such as *Drosophila* with smaller numbers of chromosomes and the lack of recombination in males (Lynch et al. 2017). Genotype frequencies in these populations typically adhere very closely to Hardy-Weinberg expectations (Maruki et al. 2022).

On the other hand, as residents of permanent lakes, *D. pulicaria* are capable of foregoing sexual reproduction for multiple years, which in some cases could extend over tens to hundreds of generations. This often leads to substantial deviations from Hardy-Weinberg expectations within loci and elevated linkage disequilibrium among loci, including extreme cases of transient near-fixation of single clones within individual populations (Lynch 1987; Allen and Lynch 2012). *D. pulicaria* typically live in stratified lakes containing zooplanktivorous fish, where they engage in a defensive behavior known as diel vertical migration, allowing them to migrate to deeper, darker waters during the day and returning to surface waters at night to feed (Gliwicz 1986; Glaholt, et al. 2016). Although ages at first reproduction are similar in the two species, lifespans in *D. pulicaria* are double those in *D. pulex* (Dudycha 2003), likely associated with the greater opportunities for longer life in lake environments.

Whereas the average level of sequence divergence between the two species is just *∼* 1% (in excess of within-species variation) (Tucker et al. 2013; Ye et al. 2019, 2021b), the opportunities for gene flow may be limited by prezygotic isolation in nature, as some evidence suggests that sexual reproduction is induced by long-day photoperiods in *D. pulex* but by short-day photoperiods in *D. pulicaria* (Deng 1997). Nonetheless, the two species are physically capable of hybridization in the laboratory (Heier and Dudycha 2009). F_1_ hybrids in nature are typically obligate asexuals (Hebert et al. 1993; Crease et al. 1990; Xu et al. 2013, 2015), implying a breakdown in meiotic processes in hybrid backgrounds and minimal opportunities for backcrossing (Ye et al. 2021a, b). In addition, an ∼ 1.2-Mb region in *D. pulex* is thought be segregating for an introgressed fragment of the *D. pulicaria* genome that confers the inability to produce male offspring (Ye et al. 2019). A recent large-scale population-genomics study suggests that a typical *D. pulex* genome contains a small (*<* 1%) fraction of discernible *D. pulicaria-* derived DNA (Maruki et al. 2022). On the other hand, analyses involving the mitochondrial genome suggest a more reticulate evolutionary pattern, with multiple introductions of *D. pulex* mtDNA into *D. pulicaria* (Cristescu et al. 2012; Ye et al. 2022).

In summary, the existing data suggest that the closely related sister taxa *D. pulex* and *D. pulicaria* experience limited gene flow despite their effectively sympatric distributions, presumably reinforced by ecological differentiation. Here, we carry out population-genomic analyses for multiple populations of the lake-dwelling *D. pulicaria,* taking advantage of prior work on *>* 800 sequenced genomes of North American *D. pulex* distributed over ten populations (Lynch et al. 2017, 2022; Maruki et al. 2022). Comparisons of the two enable us to quantify the magnitude and patterns of nucleotide diversity and recombination rates differences. In addition, data at the whole-genome level provide further insight into the degree to which genomic regions are impacted by hybridization as well as an evaluation of patterns of differences in rates of molecular evolution between the two species.

## Results

Here, we primarily report on new observations for populations of *D. pulicaria*, integrating the results with those from broad surveys recently reported on the population-genomic features of *D. pulex* (Lynch et al. 2017, 2022; Maruki et al. 2022) throughout the narrative.

### Within-population genetic variation

In total, this study encompassed the genomic sequences of 233 clones from three *D. pulicaria* populations from permanent lakes in the midwestern United States and southern Canada, 70 to 84 clones per population (Supplementary Table S1). On average, 1.2*×*10^8^ nucleotide sites were assayed per population (Table 1), 98.7% of which were monomorphic for the same allele across all populations (Supplementary Table S2). Among polymorphic sites, *>*96% contain just two nucleotides per site, i.e., tri- and tetra-allelic sites are rare, as in the case of *D. pulex* (Maruki et al. 2022). The mean genome-wide nucleotide diversity (π) is similar across these *D. pulicaria* populations, averaging 0.0026 (SE = 0.0004), which is about one-third of that within individual *D. pulex* populations (Table 1). Although the mean inbreeding coefficient in the CLO sample is positive, it is negative for the other two populations, substantially so for population BRA. This yields an overall average *F_IS_*(inbreeding coefficient) estimate that is significantly negative (excess average heterozygosity), in contrast to the situation in *D. pulex* where average *F_IS_* is close to zero. For population CLO, 17% of the polymorphic sites exhibit deviations from Hardy-Weinberg (HW) genotype-frequency expectations, 34% of which were in the direction of heterozygote excess. In contrast, for population BRA, 50% of sites deviated from HW expectations, 84% in the direction of heterozygote excess, with these numbers becoming 16% and 65%, respectively, for population TF.

**Table 1.** Summary of within-population genetic variation. πT is the mean nucleotide diversity over all sites; the standard errors of the population estimates (not shown) are all < 0.00002. The average inbreeding coefficients, *F_IS_*, are based on polymorphisms with maximum-likelihood estimates of minor-allele frequencies significantly *>* 0.02 at the 5% level. Effective population size estimates, *N_e_*, estimated as π*_s_*/4μ, were obtained from diversity at silent sites and restricted intron positions (internally, 8 to 34 from both ends; Lynch et al. 2017), using the mutation rate estimate of u = 5.69×10^−9^ per site per generation from Keith et al. (2016).

| Population | $\pi_T$ | No. of Sites | $F_{IS}$ | SE($F_{IS}$) | No. of SNPs | $N_e$ |
| --- | --- | --- | --- | --- | --- | --- |
| BRA | 0.0029 | 117,123,707 | -0.1850 | 0.0003 | 1,656,187 | 304,000 |
| CLO | 0.0017 | 120,113,119 | 0.0137 | 0.0003 | 1,034,526 | 165,000 |
| TF | 0.0031 | 117,149,728 | -0.0340 | 0.0002 | 1,877,097 | 304,000 |
| *D. pulicaria* | 0.0026 |  | -0.1259 |  |  | 258,000 |
| SE | 0.0004 |  | 0.0317 |  |  | 46,000 |
| *D. pulex* | 0.0078 |  | -0.0168 |  |  | 640,000 |
| SE | 0.0004 |  | 0.0100 |  |  | 31,000 |

Consistent differences in nucleotide diversity exist among functionally different genomic sites (Figure 1). Nucleotide diversity is highest at silent sites, slightly lower at restricted intron sites (internal positions 8 to 34 bp from both ends; Lynch et al. 2017), and lowest at amino-acid replacement sites, with intergenic sites having intermediate levels. These observations are consistent with a predominant role for purifying selection in decreasing heterozygosity estimates at functionally more important sites, as found in essentially all species, including *D. pulex* (Maruki et al. 2022).

**Figure 1.**
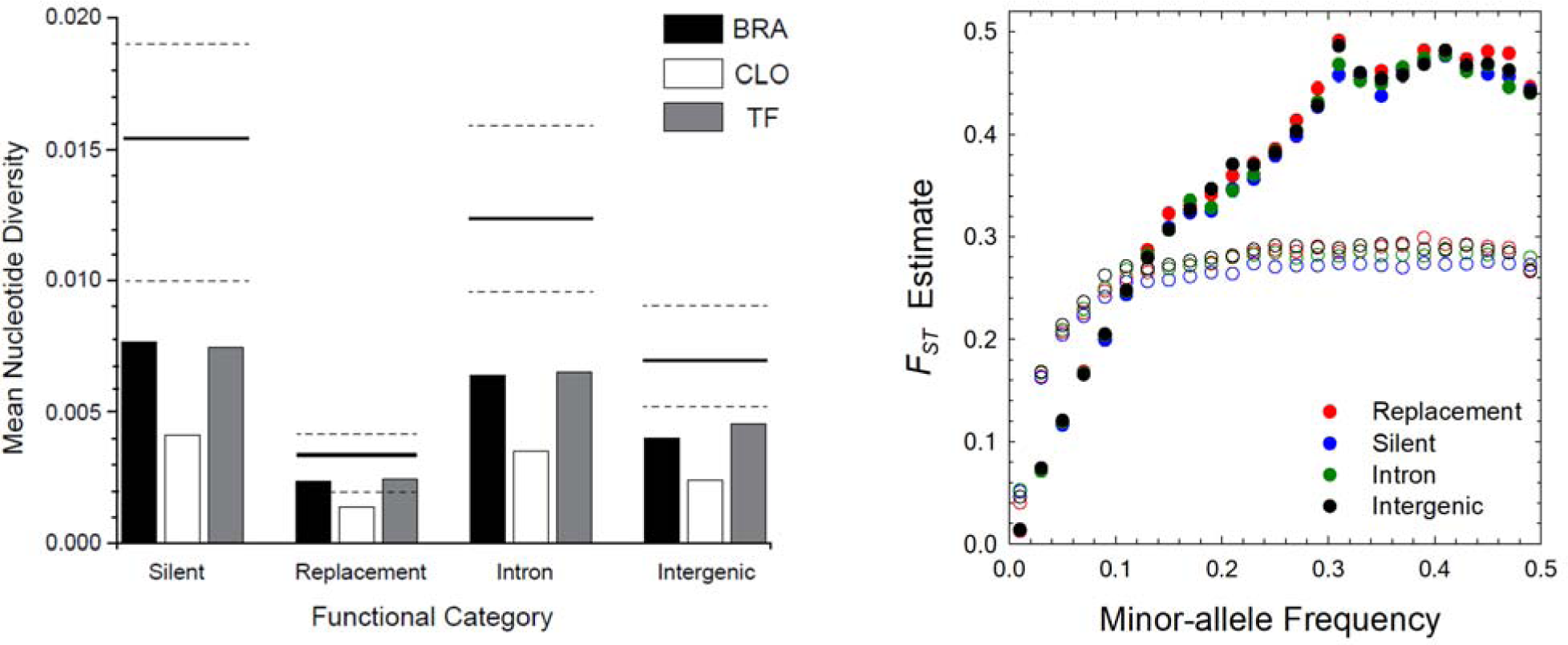
Measures of within- and among-population nucleotide variation. **Left**) Genome-wide within-population nucleotide diversity estimates at sites in different functional categories. Silent, replacement, and intron sites are, respectively, 4-fold redundant, 0-fold redundant, and restricted intron (internal position 8 to 34 from both ends; Lynch et al. 2017) sites; intergenic sites are those *>* 500 bp from translation initiation and termination codons. Tri- and tetra-allelic sites are excluded from the calculations, but make trivial contributions. The standard errors of the estimates are too small to visualize on the graph (< 10^−5^). The solid and dashed lines denote average, upper, and lower limit of estimates from *D. pulex* populations (from Maruki et al. 2022). **Right).** A comparison of *F_ST_* profiles of *D. pulicaria* (closed circles) and *D. pulex* (open circles) as a function of minor-allele frequencies at the metapopulation level.

Using the average estimates of nucleotide diversity at silent sites within codons and internal intron sites, which we have inferred to be evolving in a nearly neutral fashion (Lynch et al. 2017), and the estimated mutation rate per nucleotide site of *u* = 5.69 *×* 10^9^ per generation for *D. pulex* (Keith et al. 2016), we infer the mean effective population size of *D. pulicaria* to be ∼ 40% of that for *D. pulex* (Table 1). Similarly, Wersebe et al. (2022) estimated that *D. pulicaria* has an effective population size about a third that of *D. pulex*.

### Divergence among subpopulations

Owing to the downward bias of the *F_ST_* measure of population subdivision with low minor-allele frequencies (MAFs) (Alcala and Rosenberg 2017), estimates of this index increase with increasing MAF, leveling off at the point at which the latter exceeds ∼ 0.3, with minimal differences among different functional site types (Figure 1). The overall mean *F_ST_* (relative divergence between two populations) for *D. pulicaria* is 0.227 (SE = 0.0001), whereas focusing just on silent sites with MAF *>* 0.15, the mean increases to 0.411 (0.001), and for such sites with MAF *>* 0.30, it further increases to 0.467 (0.0003).

Given the similarity of *N_e_* estimates across populations (Table 1), and the similar behavior of all functional categories of sites, an estimate of the average migration rate using equilibrium theory appears to be justified as a first-order approximation. To this end, we used Wright’s (1951) island model wherein a metapopulation consists of a large number of demes of size *N_e_,* with each deme experiencing an equivalent migration rate of *m* per individual (randomly distributed among demes), which yields an expected equilibrium value of *F_ST_*of (1 + 4*N_e_m*)*^-^*^1^. Setting *F_ST_* = 0.467, the mean effective number of migrants per deme per generation is estimated to be *N_e_m* = 0.29 for *D. pulicaria,* which with the average *N_e_* in Table 1 implies a migration rate of *m ∼* 1.1*×*10^-6^. In contrast, the unbiased estimated mean *F_ST_* for *D. pulex* populations is 0.269 (0.002), with a mean number of migrants per generation of *N_e_m ∼* 0.59, and a migration rate of *m* = 9.2×10^−7^ (Maruki et al. 2022). Thus, both species have per-capita migration rates ≈ 10^−6^, whereas *D. pulex* has essentially double the absolute migration rate of *D. pulicaria,* owing to the elevated *N_e_.* The net effect of the latter is a substantial overall reduction in the level of population subdivision in *D. pulex*.

The elevated level of population subdivision in *D. pulicaria* is not a consequence of increased geographic distance between the sampled populations. The pairwise average *F_ST_* (MAF>0.3) for BRA vs. CLO is 0.521, for BRA vs. TF is 0.262, and for CLO vs. TF is 0.535 (all with SE ≈ 0.005). *D. pulex* exhibits a weak pattern of isolation by distance (Maruki et al. 2022), and for the distances separating the above pairs (551, 787, and 524 km, respectively), the expected *F_ST_* for *D. pulex* falls in the narrow range of 0.246 to 0.270, implying reduced migration per physical distance in *D. pulicaria*.

Finally, some idea of the degree to which the results from these three populations reflect the overall features of the entire North American metapopulation of *D. pulicaria* can be acquired by considering an evaluation of the genomic sequences of 14 clones distributed over the North American continent, from east to west coasts (Jackson et al. 2021). Because only a single clone was sampled per population in the latter study, the average within-population diversity cannot be estimated. However, the total variation across populations should provide a good estimate of the species-wide diversity, as the average within-population variance will be contained within the overall sample. The total nucleotide diversities are: 0.0134, 0.0109, 0.0077, and 0.0045 for synonymous, intron, intergenic, and nonsynonymous sites, respectively, whereas the metapopulation-wide diversities for the three populations reported herein are 0.0109, 0.0098, 0.0063, and 0.0033, respectively (in all cases, the SEs of these measures are *<* 0.0001). Thus, the entire species appears to harbor ∼ 23, 11, 22, and 36% more diversity for these categorical sites than revealed in our three study populations.

### Private alleles within populations

An alternative way to consider population subdivision evaluates the incidence of significant private alleles (SPAs), defined to be polymorphisms that are restricted to single populations and have an observed frequency that is significantly greater than expected under the null hypothesis of random sampling from a single panmictic population (Maruki et al. 2022). The maximum frequency of a private allele in a set of *n* populations is equal to *p/n,* where *p* is the metapopulation MAF, and rare alleles are unlikely to be designated as SPAs owing to the high probability of emerging by chance sampling of frequency 0.0 in all but one population. In *D. pulex* populations, the fraction of SPAs among all polymorphisms exhibits an exponential decline with increasing metapopulation MAF (Maruki et al. 2022). In striking contrast, in these *D. pulicaria* populations, the incidence of SPAs is quite nonresponsive to the metapopulation MAF (Figure 2). Between 19,000 and 40,000 SPAs were found in each population, with their incidence declining with increasing frequency in the focal population (Figure 2). Owing to the fact that ten populations of *D. pulex,* but only three of *D. pulicaria* enter these analyses, the maximum frequency of a SPA in the former is about one-third of that in the latter. Nonetheless, it remains clear that *D. pulicaria* populations have exceptionally high incidences of high-frequency SPAs (Figure 2). This condition is almost certainly a consequence of prolonged clonal selection in the absence of recombination resulting in the periodic emergence of a few clones in each population, which drags all SNPs within such clones to high frequency.

**Figure 2.**
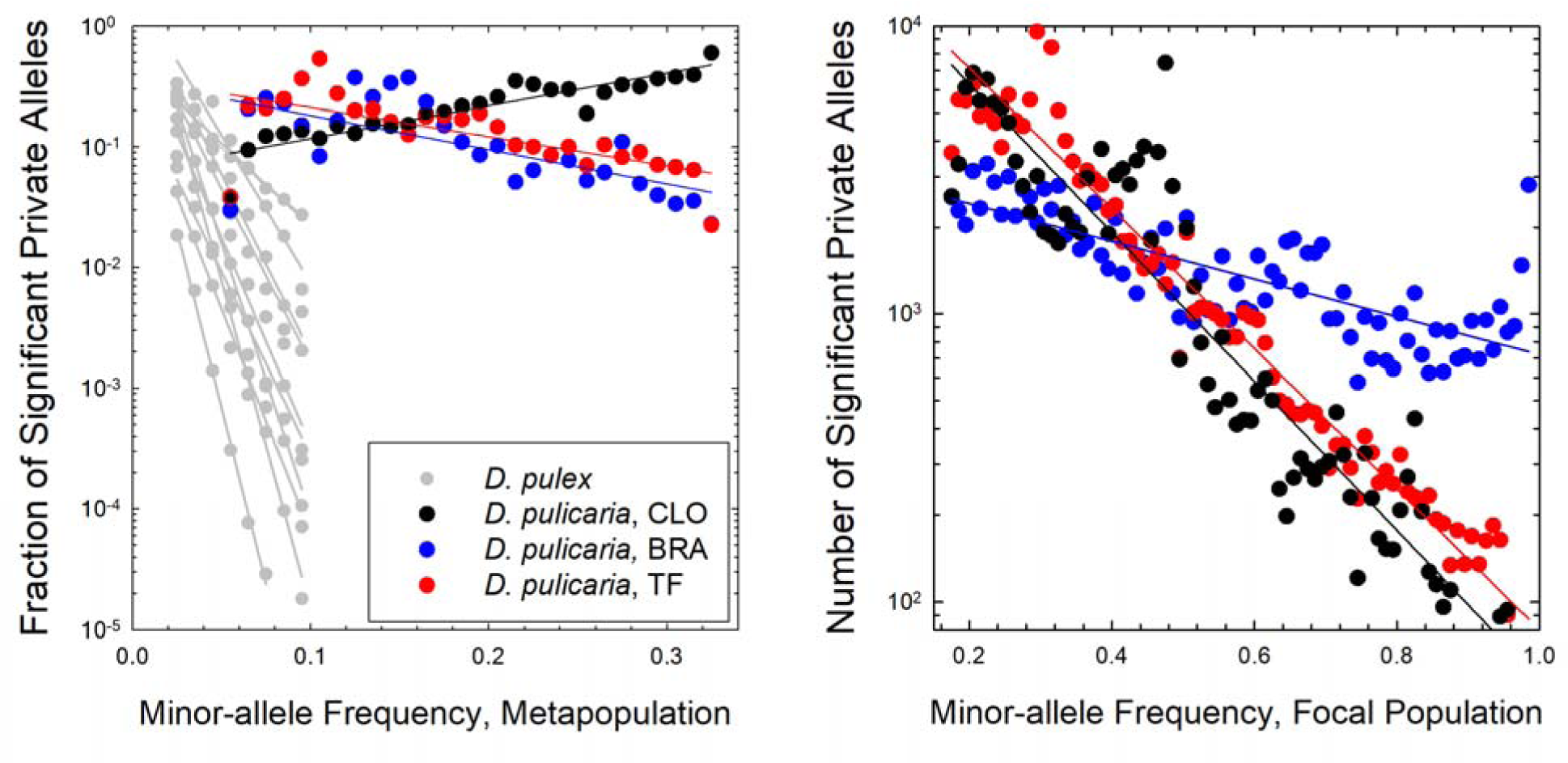
Incidence of private alleles in the three *D. pulicaria* populations. **Left**) The fraction of private SNPs with the designated metapopulation-wide MAF. Gray data points and regression lines for *D. pulex* are from Maruki et al. (2022). **Right)** Counts of significant private alleles in each focal population as a function of the local MAF.

Using the genome-wide estimate of the frequency of private alleles per site as the null hypothesis, for each protein-coding gene in each population, assuming a Poisson distribution for the number of private alleles over entire genic regions (from the start of the 5^’^ UTR to the termination of the 3^’^ UTR), we determined the subset of genes containing significantly more private alleles than expected by chance in each population (as in Maruki et al. 2022). After Bonferroni correction, this analysis revealed that 134, 112, and 80 genes in the CLO, TF, and BRA populations, respectively, were enriched for private alleles beyond the background expectation, none of which overlap among populations (Supplemental File S1). On average, these enriched genes had 13.0 (SE = 0.4) private SNPs, with a mean within-population private-allele frequency of 0.345 (0.0003). Thus, although restricted to individual *D. pulicaria* populations, these divergent haplotypes are locally quite abundant.

As noted above, the incidence of private alleles in a metapopulation sample will depend on the number of distinct population samples, declining as the latter increases. Nonetheless, several features of the *D. pulicaria* private-enriched genes are strikingly apparent. First, they are strongly enriched on chromosome 11 (29%) in BRA and chromosome 4 (30%) in TF (Figure S1). As all 12 *D. pulicaria* chromosomes are similar in physical length, this represents a highly significant elevation of incidence of private alleles on these two chromosomes (Figure S1). Second, the private-enriched genes are somewhat concentrated in the “islands of divergence” identified in Jackson et al. (2021) for this species, most of which are associated with chromosomal rearrangements with respect to *D. pulex;* 59.8% of the *D. pulicaria* private-enriched genes are located in such regions, which constitute only 36.4% of the *D. pulicaria* genome.

To evaluate whether the *D. pulicaria* private SNPs might result from introgression, we determined the fraction of such SNPs that match known nucleotide variants in the most closely related species, *D. pulex,* as the two are known to hybridize in the field and in the laboratory (Heier and Dudycha 2009; Vergilino et al. 2011; Xu et al. 2015; Moy et al. 2021). Notably, although 8.6% of *D. pulicaria* private SNPs are also found in *D. pulex,* this incidence is twice as enriched (17.3%) in coding regions. Nearly half (46.9%) of private-enriched *D. pulicaria* genes contain private SNPs matching *D. pulex* nucleotides (Maruki et al. 2022), with an average fraction 0.162 (SE = 0.006) of the private SNPs in such genes matching unique variants in *D. pulex*. Thus, there is little question that some gene sharing occurs between these two species, and that such overlap is particularly enriched in protein-coding genes, although the number of genes involved comprises *<*1% of the genomic total within each *D. pulicaria* population. About 18% of the private *D. pulicaria* SNPs are fixed in *D. pulex* metapopulations, a strong indication of gene flow from *D. pulex*. However, the majority (82%) of the private SNPs are still polymorphic in the *D. pulex* metapopulation, so we are unable to determine whether they represent shared ancestral polymorphism or result from ongoing gene flow.

### Site-frequency spectra

A site-frequency spectrum (SFS) describes the genome-wide distribution of SNP frequencies, here described by the folded SFS based on minor-allele frequencies (MAFs, with frequencies between 0.0 and 0.5), as we are not yet able to fully assign ancestral states across the genome. Under neutral evolution in a single panmictic population of constant size, the folded SFS is expected to scale as 1*/*[*p*(1-*p*)], where *p* is the minor-allele frequency, with the elevation of the SFS being defined by the composite parameter 4*N_e_u* (Wright 1938; Messer 2009). As was previously found for *D. pulex* (Maruki et al. 2022), the SFSs for four-fold redundant sites in protein-coding genes and for restricted internal intron sites are essentially indistinguishable, and these are averaged in Figure 3.

**Figure 3.**
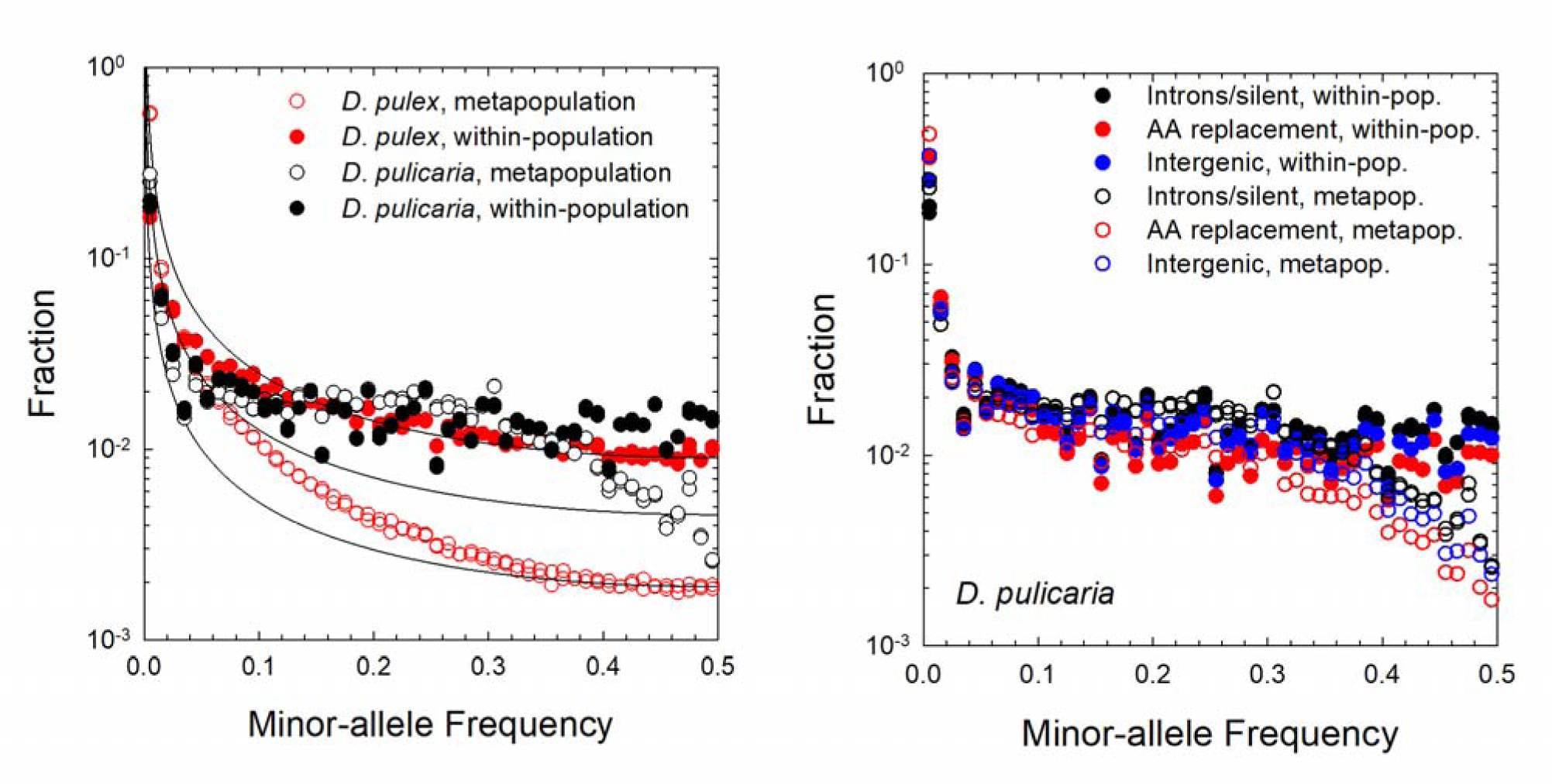
Site-frequency spectra (SFSs) for polymorphisms in *D. pulicaria* and *D. pulex.* **Left**) SFSs for putatively neutral sites given for both the average within-population pattern and for the entire metapopulation. For each MAF bin (of width 0.01), there are two points for each species, one for four-fold redundant (silent) sites in protein-coding genes and the other for restricted intron sites (8 to 34 from both ends; Lynch et al. 2017). The plotted SFSs are exclusive of the metapopulation-wide monomorphic classes. The curved lines are benchmarks of arbitrary height, showing the scaling [*p*(1 − *p*)]*^-^*^1^, where *p* is the minor-allele frequency, expected in the ideal case of neutral sites in a single panmictic, random-mating population at drift-mutation equilibrium. Data for *D. pulex* is from Maruki et al. (2022). **Right)** SFSs for functional categorical sites in *D. pulicaria* at the within-population (averaged over all three populations) and metapopulation levels.

At the average within-population level, the SFSs for neutral sites are quite similar in both species (Figure 3, left). In both cases, there tends to be a deficit of rare alleles, with the SFS otherwise corresponding fairly closely to the neutral expectation. There are, however, three substantial differences in other aspects of SFS behavior between the two species. First, whereas the metapopulation SFS for neutral sites in *D. pulex* declines in a fairly continuous manner, that for *D. pulicaria* exhibits a more discontinuous behavior. In both cases, there is an excess of rare neutral alleles, as expected when migration is limited, as young alleles require time to move and equilibrate in frequency across the metapopulation (De and Durrett 2007; Städler *et al*. 2009). However, there is a huge bulge in the neutral SFS in the MAF range of 0.1 to 0.4 for *D. pulicaria,* whereas the *D. pulex* metapopulation SFS declines in a smooth fashion. Second, the within-population SFSs for intergenic and amino-acid replacement sites are quite similar to the neutral-site SFS in *D. pulicaria* (Figure 3, right). This contrasts with the substantial depression in the frequencies of all but the lowest MAFs for these categorical sites in the *D. pulex* within-population SFS, by nearly an order of magnitude in the highest frequency sites (Lynch et al. 2017; Maruki et al. 2022). Third, the distinctions between the metapopulation SFSs for the three functional classes of sites in *D. pulicaria* remain far less extensive than observed in *D. pulex,* where all types of sites become increasingly rare with high MAFs (Maruki et al. 2022).

### Linkage disequilibrium

We measured the decline of linkage disequilibrium (LD) with physical distance along chromosomes with two different methods. First, the correlation of zygosity (Δ) is an individual-based method, based on the degree to which pairs of sites are jointly heterozygous (Lynch 2008; Lynch et al. 2014). In random-mating populations, when scaled by the average heterozygosity, Δ is expected to decline with physical distance among sites in the same manner as population-level LD. More general advantages of the approach are that it makes no assumptions about random mating and requires no haplotype phasing. Using this approach, it is clear that there is substantially more LD at all length scales in all *D. pulicaria* populations than in *D. pulex* (Figure 4A). The magnitude of inflation gradually increases from ∼ 1.5*×* at nearly adjacent sites to ∼ 5*×* at distances in excess of 10^5^ bp. These elevated levels of Δ are also seen in all of the isolates (all from different populations) evaluated in Jackson et al. (2021), showing that this is a species-wide pattern. Second, population-level LD (*r*^2^) based on genotype-frequency distributions (Rogers and Huff 2009) in *D. pulicaria* is also substantially greater than that in *D. pulex* at all length scales, exceeding 0.3 to 0.4 on average even at distances as large as 250 kb, even for low-frequency SNPs that tend to lead to downwardly biased estimates of *r*^2^ (Figure 4B).

**Figure 4.**
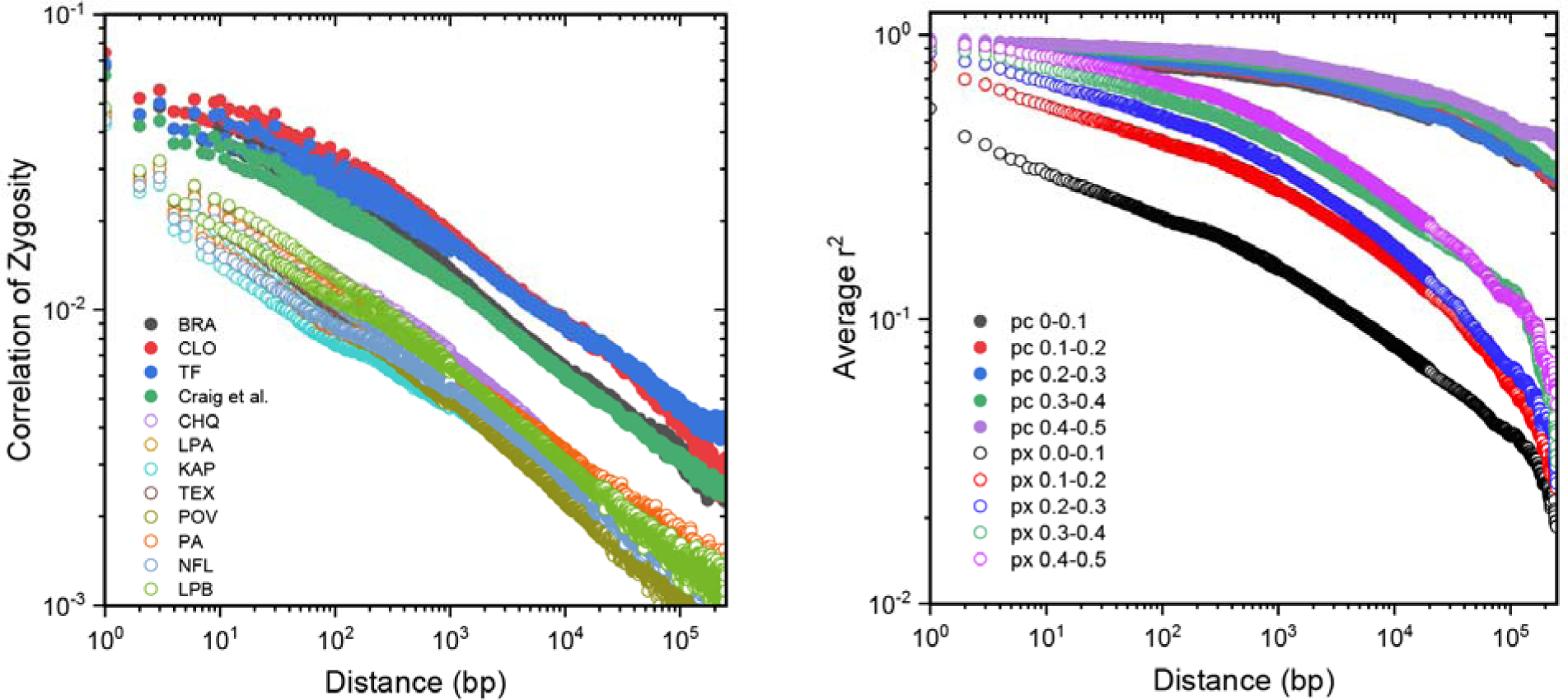
Linkage disequilibrium as a function of physical distance along chromosomes. **Left**) Individual-based correlation of heterozygosity. Craig et al. denotes points derived from a collection of single clones from 14 different populations (Jackson et al. 2022). Closed (upper) and open (lower) points are averages from surveys of ten clones from each of three populations of *D. pulicaria* and eight of *D. pulex* (Lynch et al. 2022). **Right)** Population-level measures of linkage disequilibrium given as averages over the three *D. pulicaria (pc)* populations (closed points), compared to averages from eight *D. pulex (px)* populations (open points; from Lynch et al. 2022), and given using five allele-frequency bins.

### Local adaptation associated with protein-coding genes

Given the substantial excess heterozygosity (negative *F*_IS_) in *D. pulicaria* populations, it is of interest to know whether this is reflected in unusually high levels of within-population diversity at nonsynonymous sites in protein-coding genes. However, restricting analyses to the 10,275 genes (Supplemental File S2) with π*_S_ >* 0.0005 (to avoid spurious upward bias in ratios), average π*_N_/*π*_S_* is 0.2425 (0.0034), nearly identical to the estimate for *D. pulex* obtained in the same way, 0.2504 (0.0030; 12,843 genes) (Table 3). For the subset of 9,204 protein-coding genes that could be compared across species (with confident orthology and enough data coverage), the similarity between the two taxa remains: π*_N_/*π*_S_* = 0.1961 (0.0022) for *D. pulex,* and 0.2086 (0.0026) for *D. pulicaria* (Table 3).

**Table 2.**
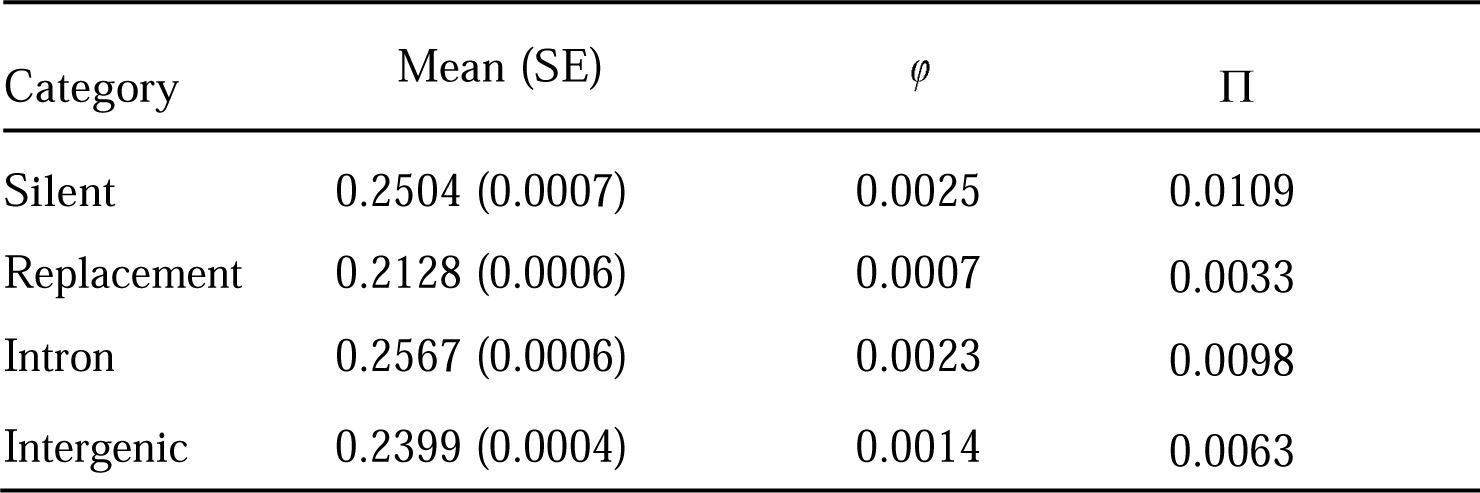
Mean *F_ST_*estimates at sites in different functional categories, along with average nucleotide diversities at the among-population (φ) and total-metapopulation (Π) levels. The means for *F_ST_* are calculated using sites with significant polymorphisms at the 5% level in at least one of the populations (SEs are in parentheses). Intergenic sites are those outside of untranslated regions (UTR), exons, and introns. Silent, replacement, and intron sites are, respectively, four-fold redundant, zero-fold redundant and restricted intron sites. Standard errors for all mean estimates of nucleotide diversity are *<* 0.00003.

| Category | Mean (SE) | $\varphi$ | $\Pi$ |
| --- | --- | --- | --- |
| Silent | 0.2504 (0.0007) | 0.0025 | 0.0109 |
| Replacement | 0.2128 (0.0006) | 0.0007 | 0.0033 |
| Intron | 0.2567 (0.0006) | 0.0023 | 0.0098 |
| Intergenic | 0.2399 (0.0004) | 0.0014 | 0.0063 |

**Table 3.** Comparison of sequence diversity and selection intensities between *D. pulex* and *D. pulicaria. All* indicates all genes with πs estimates >0.0005 (to avoid spurious upward bias in ratios). *With orthologs* indicates genes that are one-to-one orthologs between *D. pulex* and *D. pulicaria*, and *without orthologs* indicates genes lacking orthologous data in one or the other species. NI estimates are restricted to genes with NI*_B_ <* 10.0. Numbers in the parentheses are standard errors. px, *D. pulex*; pc, *D. pulicaria*.

| | | $\pi_n$ | $\pi_s$ | $\pi_n/\pi_s$ | $d_n$ | $d_s$ | $d_n/d_s$ | $NI_B$ |
| --- | --- | --- | --- | --- | --- | --- | --- | --- |
| *All* | px | 0.0029<br>(0.0000) | 0.0160<br>(0.0001) | 0.2504<br>(0.0030) | 0.0172<br>(0.0002) | 0.1232<br>(0.0005) | 0.1511<br>(0.0038) | 1.4779<br>(0.0124) |
|  | pc | 0.0015<br>(0.0000) | 0.0081<br>(0.0001) | 0.2425<br>(0.0034) | 0.0183<br>(0.0002) | 0.1303<br>(0.0005) | 0.1531<br>(0.0044) | 1.6464<br>(0.0015) |
| *With*<br>*orthologs* | px | 0.0026<br>(0.0000) | 0.0169<br>(0.0001) | 0.1961<br>(0.0022) | 0.0170<br>(0.0001) | 0.1230<br>(0.0004) | 0.1478<br>(0.0034) | 1.3796<br>(0.0125) |
|  | pc | 0.0013<br>(0.0000) | 0.0081<br>(0.0001) | 0.2086<br>(0.0026) | 0.0172<br>(0.0002) | 0.1301<br>(0.0005) | 0.1401<br>(0.0056) | 1.6147<br>(0.0171) |
| *Without*<br>*orthologs* | px | 0.0038<br>(0.0000) | 0.0141<br>(0.0002) | 0.4352<br>(0.0109) | 0.0242<br>(0.0004) | 0.1221<br>(0.0013) | 0.2591<br>(0.0122) | 1.4779<br>(0.0124) |
|  | pc | 0.0026<br>(0.0001) | 0.0079<br>(0.0003) | 0.5359<br>(0.0066) | 0.0297<br>(0.0009) | 0.1359<br>(0.0006) | 0.2685<br>(0.0035) | 1.7826<br>(0.0169) |

To identify genes potentially under diversifying selection at the amino-acid level, we tested the difference between π*_N_* and π*_S_* in units of sampling standard errors. Of the 11,205 *D. pulicaria* genes that could be evaluated, 733 have π*_N_ /*π*_S_* values exceeding 1.0, but with a critical Z-score value of 4.46 after correcting for multiple comparisons, only 51 (0.45%) are significant. This is more than double the incidence of such significant outlier genes in *D. pulex* (0.22%; Maruki et al. 2022). However, 15 of these 51 genes are not found in the annotated *D. pulex* genome (including six out of the ten genes with the highest π*_N_ /*π*_S_* values, with values ranging from 14.8 to 53.7), and of the remaining 36, only five have π*_N_ /*π*_S_ >* 1.0 in *D. pulex*. Notably, only two of these genes have π*_N_* greater than the average π*_S_* for all *D. pulicaria* genes, 0.0081 (0.0001), the average π*_S_* for the 51 being just 0.0011 (0.0003). Thus, it is plausible that the majority of these high π*_N_ /*π*_S_* genes are stochastic results of low π*_S_* rather than indicators of elevated π*_N_,* although the latter has an average of 0.0033 (0.0007), which is ∼ 3*×* higher than the genome-wide mean of 0.0015 (0.00002). We tentatively conclude that the incidence of balancing selection at amino-acid replacement sites in these populations is very low. Interestingly, for genes without orthologs across both species, i.e., species-specific genes, π*_N_* exhibited significant elevations in both species, although π*_S_* remains similar to genes with orthologs (Table 3). These results suggest that, at the amino-acid sequence level, newly generated genes in each lineage may evolve faster than genes with shared ancestry.

To determine whether there are any genes under strong diversifying selection among the three populations, we considered the ratio φ*_N_ /*φ*_S_,* where φ*_x_* is the among-deme measure of nucleotide diversity. This ratio is expected to exceed 1.0 in cases of strong local adaptation at the amino-acid sequence level. Of the 10,406 genes that could be evaluated (Supplemental File S2), only in 48 cases (*<* 0.5%) was φ*_N_* more than two standard errors greater than φ*_S_,* and after accounting for multiple comparisons, none of these retained significance. With only three populations evaluated, this analysis does not have great power, but it is notable that two prior studies, involving *D. pulex* (Maruki et al. 2022) and *D. magna* (Fields et al. 2015), also failed to find evidence of local adaptation with this approach.

As tests for π*_N_ /*π*_S_* > 1 and φ*_N_ /*φ*_S_ >* 1 are quite stringent, owing to the likelihood that a substantial fraction of mutations is deleterious even on a background of positive selection, in a further search for genes exhibiting a signature of local adaptation, we considered the metapopulation neutrality index, NI*_W_*, which is the ratio of π*_N_ /*π*_S_* to φ*_N_ /*φ*_S_* (Maruki et al. 2022). Protein-coding genes under diversifying selection among demes are expected to have excess amino-acid sequence divergence (scaled to the silent-site diversities), i.e., NI*_W_ <* 1. For the 6,376 genes that could be compared across both species, the average index is 2.27 (0.09) in *D. pulex* and 3.66 (0.34) in *D. pulicaria.* Removing outliers with NI*_W_ >* 10, which constitute 2.6% of genes in *D. pulex* and 5.1% of genes in *D. pulicaria,* these means become nearly identical, 1.56 (0.01) and 1.55 (0.02), respectively. The medians and modes of the distributions are also quite similar between species, 1.28 and 1.10 for *D. pulex,* and 1.15 and 1.12 for *D. pulicaria.* Combined with prior results for the sister taxon *D. pulex* and together comprising nearly 1000 sequenced genomes, these three sets of analyses for individual protein-coding genes provide little evidence for diversifying / balancing selection within or among populations.

### Between-species divergence

To obtain insight into longer-term aspects of evolution of protein-coding genes, we considered levels of divergence with respect to the outgroup *D. obtusa.* The full set of 10,439 genes with adequate data for at least two *D. pulicaria* populations and putative orthologs in *D. obtusa* yielded average values of *d_N_* = 0.0183 (0.0002)*, d_S_* = 0.1303 (0.0005), and *d_N_ /d_S_* = 0.1531 (0.0044) (Table 3). Comparable average values for the full set of 9,599 *D. pulex / D. obtusa* orthologs are: *d_N_* = 0.0172 (0.0002)*, d_S_* = 0.1232 (0.0005), and *d_N_ /d_S_* = 0.1511 (0.0038) (Table 3). Considering just the 9,467 protein-coding genes for which both species could be compared to the outgroup, mean *d_N_* is essentially identical for both species: 0.0172 (0.0002) for *D. pulicaria,* and 0.0170 (0.0001) for *D. pulex* (Table 3). On the other hand, mean *d_S_*is slightly higher for *D. pulicaria* than for *D. pulex,* 0.1301 (0.0005) and 0.1230 (0.0004), respectively (Table 3). This then renders mean *d_N_ /d_S_* slightly lower for *D. pulicaria* than for *D. pulex* orthologs, 0.1401 (0.0056) and 0.1478 (0.0034), respectively (Table 3). For genes without orthologs in either species, *d_N_* exhibited significant elevations compared to genes with *D. pulex/pulicaria* orthologs, leading to higher *d_N_/d_S_*values (Table 3).

Using the three two-species divergence estimates, the overall levels of divergence on the *D. pulex* and *D. pulicaria* branches to their common ancestor (rooted by *D. obtusa*) can be obtained by the solution of simultaneous equations (Sarich and Wilson 1967; Wu and Li 1985). This reveals a 29% increase in the amount of divergence at silent sites on the *D. pulicaria* vs. the *D. pulex* branch. This need not imply a difference in mutation rates between taxa, however, as there are almost certainly differences in annual numbers of generations between the two. Were there an average of 29% more generations/year in *D. pulicaria* than in *D. pulex* populations, for example, the above results would imply equal per-generation mutation rates in the two taxa. Given that *D. pulicaria* occupies permanent lakes, whereas *D. pulex* is dormant most of the year, it is plausible that *D. pulicaria* has twice the number of annual generations, in which case the difference in silent-site divergence would imply a *D. pulicaria* mutation rate ∼64% of that in *D. pulex*.

To gain further insight into whether genes on the *D. pulicaria* vs. *D. pulex* branches experience different levels of purifying and/or positive selection relative to the outgroup, we evaluated the between-species neutrality index NI*_B_*, which is the ratio of Π*_N_ /*Π*_S_* to *d_N_ /d_S_*, where Π*_x_* = π*_x_* + φ*_x_* is the total nucleotide diversity in the metapopulation of a species. The expected value of NI*_B_* is equal to 1.0 under neutrality, declines towards 0.0 as positive selection promotes amino-acid sequence divergence between species, and exceeds 1.0 when the majority of amino-acid substitution sites are under purifying selection. Regardless of the measure of statistical centrality, these analyses suggest higher NI*_B_*for *D. pulicaria* than for *D. pulex.* Excluding genes with extreme values >100.0, mean NI*_B_* = 3.42 (SE = 0.09; N = 10,197) in *D. pulicaria* and 2.10 (SE = 0.05; N = 9,506) in *D. pulex.* Further restricting the analyses to genes with NI*_B_ <* 10.0, the respective values become 1.65 (SE = 0.02; N = 9,550) and 1.48 (SE = 0.01; N = 9,244). Using the full set of genes for each species, the respective medians are 1.26 vs. 1.12, and the respective modes are 0.94 vs. 0.86. Whereas the among-gene variance in NI*_B_* estimates is greater for *D. pulicaria* than for *D. pulex,* this may largely be a consequence of the former being based on three populations and the latter based on ten. These results also remain virtually identical if the analyses are restricted to *D. pulicaria / pulex* orthologs. Although the tendency for most of these centrality measures to exceed 1.0 implies an overall predominance of purifying selection acting on amino acid altering mutations, mode values *<* 1 imply some operation of positive selection with respect to divergence, more so in *D. pulex*.

For the 9,858 *D. pulicaria* genes that could be tested (with confident orthology with *D. obtusa* and enough data coverage for at least two populations), 1,578 had NI*_B_* estimates significantly smaller than 1.0 (using critical Z scores at the 5% level) before Bonferroni correction, and with a critical Z score of 4.44 after correction, 808 (8.2%) remain significant (Supplemental File S3). By comparison, 5.3% of the genes in *D. pulex* were deemed to be under significant positive selection (Maruki et al. 2022). Of these 808 *D. pulicaria* genes, 741 had parallel data for *D. pulex,* 192 (26%) of which were also flagged as being under significant positive selection in this species. Functional-enrichment analysis reveals that positively selected genes in *D. pulicaria* are concentrated in ageing and metabolic related categories such as cellular homeostasis, single-organism catabolic process, carboxylic-acid metabolic process, and small-molecule metabolic process (Figure 5), none of which are highlighted in *D. pulex* (Figure S2). In contrast, GO categories such as visual perception, one of the most significant GO categories in *D. pulex* (Figure S2), are not enriched in *D. pulicaria*. However, genes in GO categories involving ribosomes, mitochondria, and responses to environmental stimuli were found to be under positive selection in both species (Figure 5 and Figure S2).

**Figure 5.**
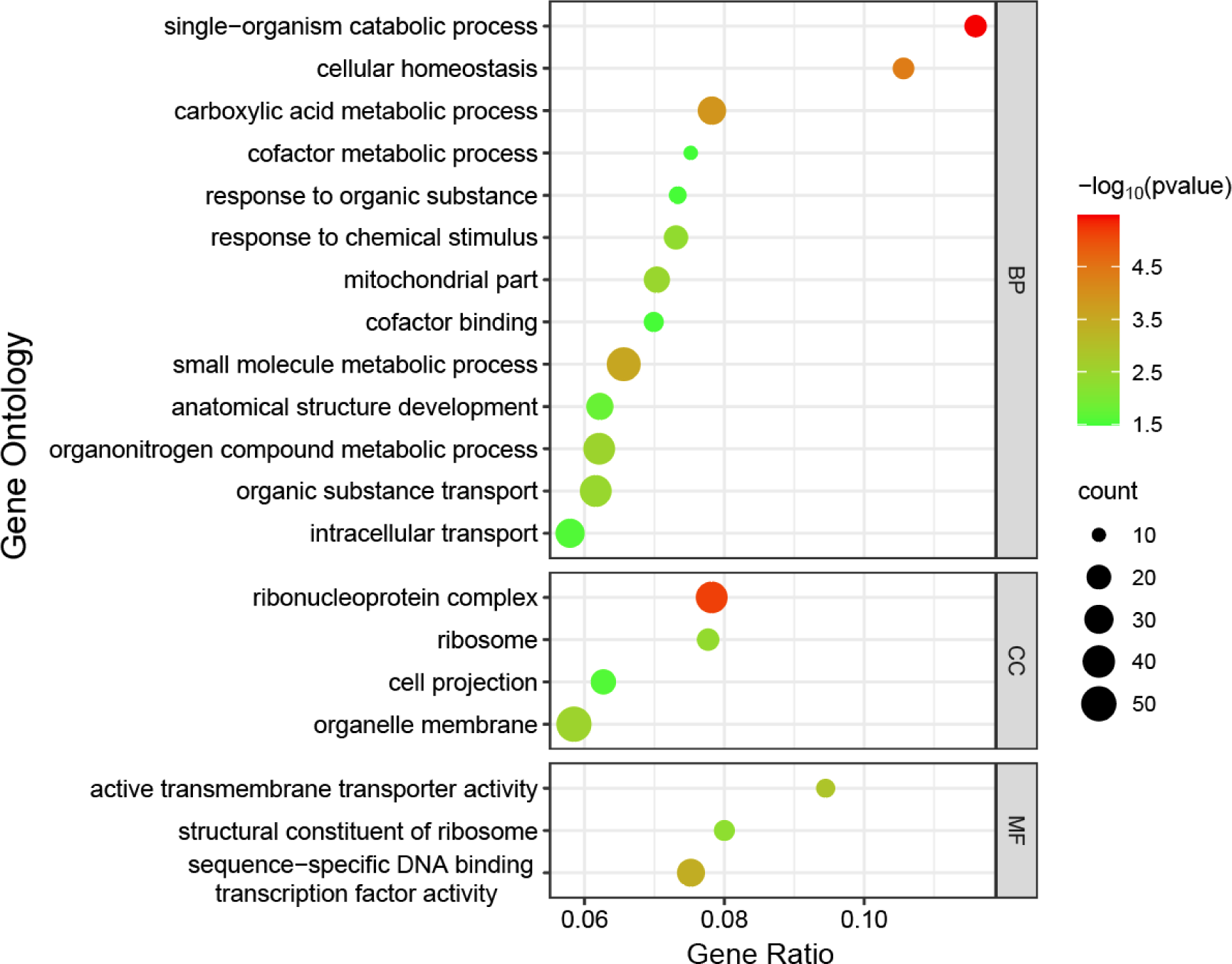
Gene-ontology (GO) categories enriched with positively selected genes in *D. pulicaria. P* values are derived from χ^2^ tests corrected for multiple testing using the Benjamini and Hochberg (1995) method. The gene ratio is the number of positively selected genes divided by the total number of genes tested under each Gene Ontology term. BP, biological process; CC, cell component; MF, molecular function.

## Discussion

Contrasting sister species experiencing ecological differentiation is of particular interest, as a central goal of evolutionary biology is to understand how species adapt to distinct habitats. Here, we compared population-genomic data between the pond-dwelling *D. pulex* and lake-dwelling *D. pulicaria*, showing that the population-genetic structure differs substantially between the two species. Genome-wide nucleotide diversity for *D. pulicaria* is much lower than that in *D. pulex*, likely reflecting a smaller *N*_e_ than in *D. pulex* (assuming equal mutation rates in the two species), despite the fact that individual populations of the former occupy much larger bodies of water. Our estimate for *N*e in *D. pulicaria* populations is very similar to that in Wersebe et al. (2022) (258,000 vs. 221,377), although the two studies are based on different populations. The latter were estimated from multiple *D. pulicaria* populations from a much wider geographic distribution (Wersebe et al. 2022). Therefore, the reduced *N*e in *D. pulicaria* compared to *D. pulex* is unlikely caused by sampling bias. Unlike *D. pulex*, *D. pulicaria* commonly exhibits heterozygosity at nucleotide sites in excess of Hardy-Weinberg expectations, as previously observed in other permanent-lake populations, including *D. pulicaria* (Hebert 1974; Morgan et al. 2001). The per-capita migration rates of the two species are fairly similar, but owing to the larger *N*_e_, *D. pulex* has a higher absolute migration rate, and hence a lower level of population subdivision. *D. pulicaria* populations also have substantially more LD at all length scales than do *D. pulex*.

At the average within-population level, the SFSs for neutral sites are similar between the two species, with a deficit of rare alleles and a close correspondence to the neutral expectation. However, there are three substantial differences in SFS behavior between the two species. First, the metapopulation SFS for neutral sites in *D. pulicaria* exhibits a more discontinuous behavior compared to *D. pulex*. Second, the within-population SFSs for intergenic and amino-acid replacement sites are quite similar to the neutral-site SFS in *D. pulicaria*, in contrast to the substantial depression observed in *D. pulex*. Third, the distinctions between the metapopulation SFSs for the three functional classes (intergenic, intron/silent, replacement) of sites in *D. pulicaria* remain far less pronounced than in *D. pulex*. These results suggest that there are significant differences in the evolutionary forces shaping the genetic diversity of these two species, with *D. pulicaria* exhibiting a more complex and diverse SFS compared to *D. pulex*. This might be related to the hitch-hiking and Hill-Robertson (selective interference) effects enhanced by elevated LD in *D. pulicaria*, which in turn is a result of a reduction in the recombination rate associated with longer periods of uninterrupted clonal propagation in this species. Moreover, Wersebe et al. (2022) found that *D. pulex* and *D. pulciaria* have experienced different levels of population expansion, hence the diverse SFS could be a joint effect of demographic changes and selection.

To determine how the difference in the evolutionary forces / ecological settings might lead to divergent phenotypes (e.g., longer lifespan, vertical migration in *D. pulciaria*; different responses to short-day photoperiods) in the two species, we searched for genes under positive selection in *D. pulicaria*. 808 genes were deemed to be under significant positive selection, 192 (26%) of which overlapped with those behaving similarly in *D. pulex*. Functional-enrichment analysis revealed that positively selected genes in *D. pulicaria* are enriched in aging-related categories such as cellular homeostasis and carboxylic acid metabolic process (Figure 5). Cellular homeostasis is known to be related to aging (Zhang and Cuervo 2008; Morimoto and Cuervo 2009; Hartl 2016; Pomatto and Davies 2017). The expression of genes in carboxylic acid metabolic processes are affected by food availability (i.e., reduced calories) (Zhang, et al. 2011), which might indirectly link to aging, as caloric restriction affects aging in numerous organisms, including *Daphnia* (Cao et al. 2001; Piper and Partridge 2007; Latta et al. 2011). Thus, it is plausible that a number of positively selected genes associated with these processes might be playing important roles in the extended lifespan in *D. pulicaria* (Dudycha and Tessier 1999; Dudycha 2003). Both this study and Wersebe et al. (2022) indicated that metabolic related genes are under different selection pressures in the species pair of *D. pulex* and *D. pulicaria*, which is likely related to various aspects of life-history such as lifespan, migration behavior, predator response, etc.

Another interesting discovery is that genes related to visual perception and cellular response to light stimuli are exclusively under positive selection in *D. pulex* (Maruki et al. 2022; Ye et al. 2023). Among the genes under selection are opsin, rhodopsin, and melanopsin, all of which have been shown to mediate photoperiodic responses (Foster et al. 1985; Collantes-Alegre et al. 2018; Zhao et al. 2018; Frau et al. 2022; Senthilan, et al. 2019). Consistently, opsin genes were also found to be under positive selection in a previous study (Jackson et al. 2021), as revealed by high *F*_st_ between *D. pulex* and *D. pulciaria. Daphnia* living in ponds and lakes have significantly different responses to photoperiod (Deng 1997), likely as a consequence of differences in ambient light duration and intensity. Thus, our results suggest that the different light responses in the two species may be mediated through evolutionary changes in opsin, rhodopsin, and/or melanopsin related genes.

## Methods

### Sample preparation and sequencing

We collected *Daphnia pulicaria* isolates from three randomly chosen permanent lakes in the midwestern United States and southern Canada in spring of 2015 and 2018 (Supplemental Table S1). As in the *D. pulex* populations (Lynch et al. 2017; Maruki et al. 2022), adult individuals were collected early in the growing season, although in this case, there is no guarantee that isolates were derived from the recent hatching of resting eggs as opposed to having been propagating by overwintering individuals. As in prior work (Maruki et al. 2022), after clonal expansion of individual isolates in the laboratory, DNA was extracted for 75 to 95 isolates per population for library preparation using the Nextera kit. The DNA from each isolate was tagged by unique oligomer barcodes, with paired-end short (150 bp) reads obtained with Illumina technology.

### Data preparation

From the FASTQ files of sequence reads, we prepared nucleotide-read quartets (counts of A, C, G, and T) necessary for the population-genomic analyses, taking various filtering steps to process high-quality data. After trimming adapter sequences from the reads with Trimmomatic (version 0.36) (Bolger et al. 2014), the reads were mapped to the *D. pulicaria* reference genome (LK16) (Jackson et al. 2021), available at the NCBI GenBank (accession number GCA_017493165.1). Novoalign (version 3.02.11) vocraft.com/) was applied with the “-r None” option to exclude reads mapping to more than one location. The SAM files of the mapped reads were converted to BAM files using Samtools (version 1.3.1) (Li et al. 2009). Marked duplicates and locally realigned indels in the sequence reads were stored in the BAM files using GATK (version 3.4-0) (McKenna et al. 2010; DePristo et al. 2011; Van der Auwera et al. 2013) and Picard. In addition, we clipped overlapping read pairs by applying BamUtil (version 1.0.13) and made mpileup files from the processed BAM files using Samtools. The final file of nucleotide-read quartets for all individuals, called a pro file, was obtained using the proview command of MAPGD (version 0.4.26) ().

Even after all of the above filtering steps, numerous sources of errors may remain in population-genomic data sets. For example, poor fits of data at a site can result when some sequence reads mapping to a site are actually derived from paralogs not found in the genome assembly. Thus, to minimize the use of potentially mismapped reads, we eliminated sites with nonbinomial deviations in read distributions using a goodness-of-fit test (Ackerman et al. 2017), when the latter identified ≥ 4 deviant individuals. To further refine the data, we removed also clones with mean coverage over sites *<* 3×, and clones with a sum of the goodness-of-fit values across the genome *<* -0.4, which can happen, for example, if a clone is contaminated with another clone. Prior to final analyses, we further refined the data by removing sites associated with putatively repetitive regions identified by RepeatMasker (version 4.0.5) (www.repeatmasker.org) using the library (Jurka et al. 2005) made on August 7, 2022. In addition, we set population-coverage (sum of depths of coverage over the clones) cut-offs to avoid analyzing problematic sites (Supplemental Table S1). After the completion of all of these filtering steps, allele and genotype frequencies in each population were estimated with the maximum-likelihood (ML) method of Maruki and Lynch (2015) using the allele command of MAPGD, and as a final precaution, sites with ML error-rate estimates *>* 0.01 were removed.

### Analyses of genetic differentiation among populations

To quantify the degree of genetic differentiation among populations, we estimated Wright’s *F_ST_* from genotype-frequency estimates using Weir and Cockerham’s method (1984), taking a procedure described in Maruki *et al*. 2022. As in Maruki *et al*. (2022), we first examined the relationship between *F_ST_*and metapopulation-wide minor-allele frequency (MAF) estimates to consider the confounding effect of the maximum possible value of *F_ST_* limited by MAF (Maruki *et al*. 2012; Alcala and Rosenberg 2017) on the analyses of the *F_ST_* estimates. The method of Weir and Cockerham was designed for estimating fixation indices from accurate genotypes with no missing data. However, for data generated by high-throughput sequencing, depths of coverage vary among sites, individuals, and chromosomes within diploid individuals, so adjustments are needed to account for variability of sample sizes. We did so by estimating the effective number of sampled individuals (*n_ei_*) for which both chromosomes are sequenced at least once (Maruki and Lynch 2015) at each site within each deme. For deme *i,*

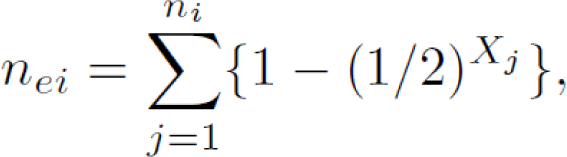

where *n_ei_* and *X_j_* are the number of sampled individuals in deme *i* and depth of coverage in individual *j*, respectively. We required that *n_ei_* be at least 10 in each deme. Letting *r_e_* denote the number of demes that satisfied this condition, the fixation indices were estimated by substituting *n_ei_* and *r_e_*for the numbers of sampled individuals in deme *i* and the number of demes, respectively, in the Weir and Cockerham equations.

### Linkage-disequilibrium analysis

To examine how linkage disequilibrium (LD) decays with distance between sites, we estimated the correlation of zygosity Δ (Lynch 2008) from the nucleotide-read quartets of single isolates using mlRho (version 2.9) (Haubold *et al*. 2010). We applied mlRho to three isolates with the highest coverage and calculated the mean of the Δ estimates over the isolates in each population. Population-level LD (*r*^2^) in *D. pulicaria* is calculated using the method in Rogers and Huff (2009), restricting analysis to significant polymorphic sites and sites with no less than 50 genotypes.

### Analyses of private alleles

To analyze alleles uniquely found in each of the populations (private alleles; Slatkin 1985), we identified private alleles taking the procedure described in Maruki *et al*. (2022). Using the mean frequencies of the private alleles over sites, the migration rate was inferred in each population using Equation 3 in Slatkin (1985) and adjusting the sample size. To infer the contribution of introgression from closely related species to the observed private alleles in genes significantly (*p <* 0.05) enriched with private alleles, we aligned reference sequences of *D. pulex* (KAP4) and *D. obtusa* to LK16 genome assembly using LAST (version 1066) and examined the fractions of private alleles matching reference nucleotides of *D. pulex* (KAP4), *D. obtusa*, and both species in each of the genes.

### Analyses of selection signatures in protein-coding sequences

To identify signatures of natural selection associated with functional changes, we compared per-site differences in protein-coding sequences between amino-acid replacement and silent sites at three levels (between species, among populations, and within populations), as in Maruki *et al*. 2022. To estimate divergence between species, we mapped sequence reads of the *D. obtusa* draft reference sequence to LK16 and called genotypes using HGC (Maruki and Lynch 2017) setting the minimum and maximum coverage cut-off values at 6 and 100, respectively, and requiring the error-rate estimate ≤ 0.01. Then, we converted homozygous genotype calls into nucleotides. Using the estimates of the divergence and heterozygosity within populations, we also estimated analogs of the neutrality index (NI) (Rand and Kann 1996). The analysis of among-population heterozygosity φ estimates was carried out using φ calculated as Π *−* π, where Π and π are the heterozygosity in the total population and mean heterozygosity within populations, respectively, and estimated in the framework of Weir and Cockerham (1984).

### Analysis of selection on coding-region DNA

To infer the form of natural selection operating on the amino-acid sequences of protein-coding genes, we estimated diversity at synonymous and nonsynonymous sites, π*_S_* and π, for each gene within each population, as well as the ratio π*_N_ /*π*_S_*. For any particular site in a particular population, π is simply estimated as 2*p*(1 − *p*),where *p* is the minor-allele frequency estimate. To minimize spurious behavior of ratios of estimates, the final gene-specific estimates of π*_N_ /*π*_S_* were obtained using metapopulation-wide means in the numerator and denominator. The sampling variances of gene-specific mean π*_S_* and π*_N_* were obtained by dividing the variance of population-specific estimates by the number of populations, and the sampling variance of π*_N_ /*π*_S_* was then obtained by use of the Delta-method equation for the variance of a ratio (A1.19b, in Lynch and Walsh 1998). In addition, the framework of Weir and Cockerham (1984) provided the difference between diversity estimates at the metapopulation level and the averages within populations, yielding estimates of among-population diversities for each site in each gene, which we denote as φ*_N_* and φ*_S_*. Gene-specific estimates of φ*_N_* and φ*_S_* estimates were calculated by averaging over all sites within genes, and their standard errors were again estimated with the Delta-method formulation. To estimate between-species divergence, we estimated for each protein-coding gene *d_N_* and *d_S_*, respectively the mean numbers of nucleotide substitutions per nonsynonymous and synonymous sites, by drawing comparisons with orthologous genes contained within the closely related outgroup species *Daphnia obtusa.* We mapped sequence reads of *D. obtusa* to *D. pulicaria* LK16 using BWA-MEM (version 0.7.17) (https://arxiv.org/abs/1303.3997) and called genotypes using HGC (Maruki and Lynch 2017), setting the minimum and maximum coverage cut-off values at 6 and 100, respectively, and requiring the error-rate estimate ≤ 0.01. To avoid analyzing sites with mismapped nucleotide reads, we again excluded sites involved in putatively repetitive regions identified by RepeatMasker, and also made further restrictions on the use of *D. pulicaria* genome sites as noted above. Codons containing undetermined nucleotides, gaps, and more than one nucleotide difference between species and stop codons were removed from the alignments. *d_N_*and *d_S_* estimation was then restricted to genes with alignments consisting of at least 300 sites, using an algorithm that calculates potential numbers of replacement and silent sites (Hedrick 2005) in units of codons weighted by allele frequencies in the *D. pulex* population, yielding divergence estimates similar to those by PAML (Yang 2007). For each gene, *d_N_* and *d_S_* were estimated as the mean divergence estimates across all populations, applying Jukes and Cantor’s (1969) method. Then, gene-specific *d_N_ /d_S_* estimates were obtained as the ratio of mean *d_N_* and *d_S_* estimates over populations, and the variance of the ratio was again calculated using the delta method.

### Enrichment analysis of gene ontology terms in the outlier genes

To infer the functions of outlier genes, we found gene ontology (GO) terms (Ashburner et al. 2000) associated with each of the genes in *D. pulicaria* LK16, using a procedure analogous to that described in Ye et al. (2017). To find GO terms enriched in each type of outlier genes, we carried out enrichment analysis of the GO terms by a χ2 test using the Perl module Statistics::ChisqIndep (https://metacpan.org/pod/Statistics::ChisqIndep). To reduce the redundancy among GO terms enriched in outlier genes, we removed GO terms when at least 80% of the associated genes are also found to be associated with another GO term with a larger number of associated genes.

## Data availability

The FASTQ files of the raw sequencing data for the *D. pulicaria* populations are available at the NCBI Sequence Read Archive (accession numbers SRS8677121, SRS8677125, and SRS8677128). The FASTQ files of the *D. obtusa* sequence reads are available at the NCBI SRA (accession number SAMN12816670), and the *D. obtusa* genome assembly is available at the NCBI GenBank (accession number JAACYE000000000). The *D. pulicaria* genome assembly LK16 is available at GenBank under accession GCA_017493165.1 and the annotation is available at https://osf.io/nr7t6/. The *D. pulex* KAP4 assembly and annotation can be accessed at NCBI under accession GCF_021134715.1. C++ programs for analyzing population structure (PSA) and finding private alleles (FPA) are available at https://github.com/Takahiro-Maruki/PSA and https://github.com/Takahiro-Maruki/FPA, respectively. The input files for these programs can be made using GFE v3.0, which is available at https://github.com/Takahiro-Maruki/Package-GFE.

## Acknowledgments

This work was financially supported by NIH grant R35-GM122566-01 to M. L. and NSF Enabling Discovery through GEnomics (EDGE) grant IOS-1922914 to M. L. and Andrew Zelhof (Indiana University). We thank Bret Coggins for assistance with the sample collections and Joe Shaw for sharing population BRA.

