## Supplementary Figures and Tables for "Evolutionary Genomics of Sister Species Differing in Effective Population Sizes and Recombination Rates"

**Table S1.** Information on the analyzed populations. Sample size denotes the number of analyzed isolates. The minimum and maximum (in square brackets) and mean total population coverage (the sum of the depths of coverage over the isolates) over all analyzed sites are provided in the final column.

| Population | (Latitude, Longitude) | Location | Sample Size | Population Coverage |
| --- | --- | --- | --- | --- |
| Brandy Lake (BRA) | (45.1097, -79.5236) | Ontario, Canada | 79 | [350, 4000], 1161 |
| Cloverdale Lake (CLO) | (42.5426, -85.3954) | Michigan, USA | 70 | [350, 4000], 1757 |
| Tenderfoot Lake (TF) | (46.2213, -89.5282) | Wisconsin, USA | 84 | [350, 4000], 1247 |

**Table S2.** Distribution of the number of distinct nucleotides detected per site in each population. In general, tri- and tetra-allelic sites are rare.

| Number of Nucleotides per Site | | | |
| --- | --- | --- | --- |
| Population | 1 | 2 | 3 or 4 |
| BRA | 0.98537 | 0.01414 | 0.00049 |
| CLO | 0.99102 | 0.00861 | 0.00036 |
| TF | 0.98355 | 0.01602 | 0.00043 |

**Figure S1.** Distribution of the *D. pulicaria* private-enriched genes for: A) all populations, B) BRA, C) CLO, and D) TF. Color shows the abundance of the genes.

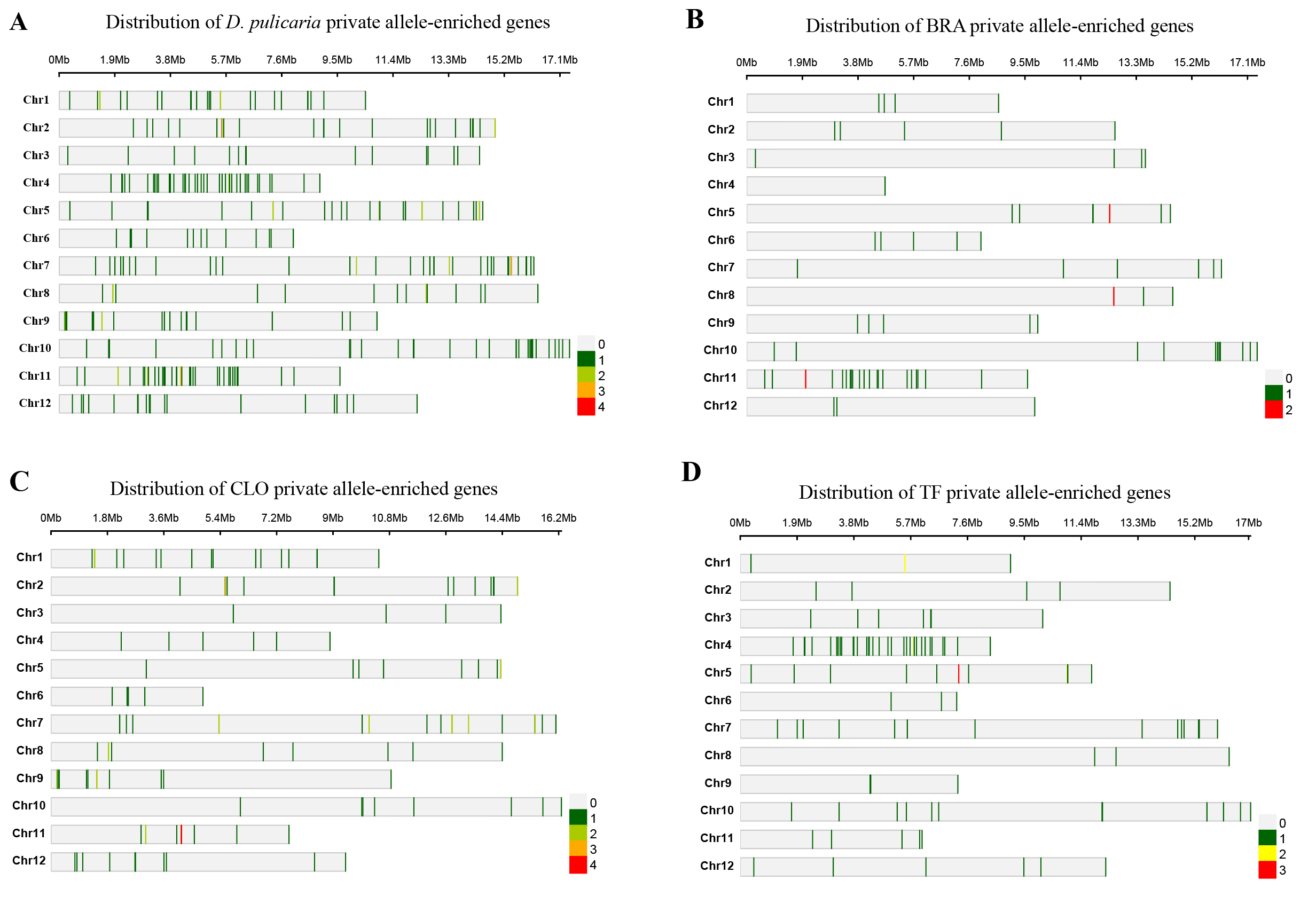

**Figure S2.** Gene ontology (GO) categories enriched with positively selected genes in *D. pulex* (Maruki et al. 2022). Ratio is calculated as the number of positively selected divided by the total tested gene in each Gene Ontology term. BP, biological process; CC, cell component; MF, molecular function.

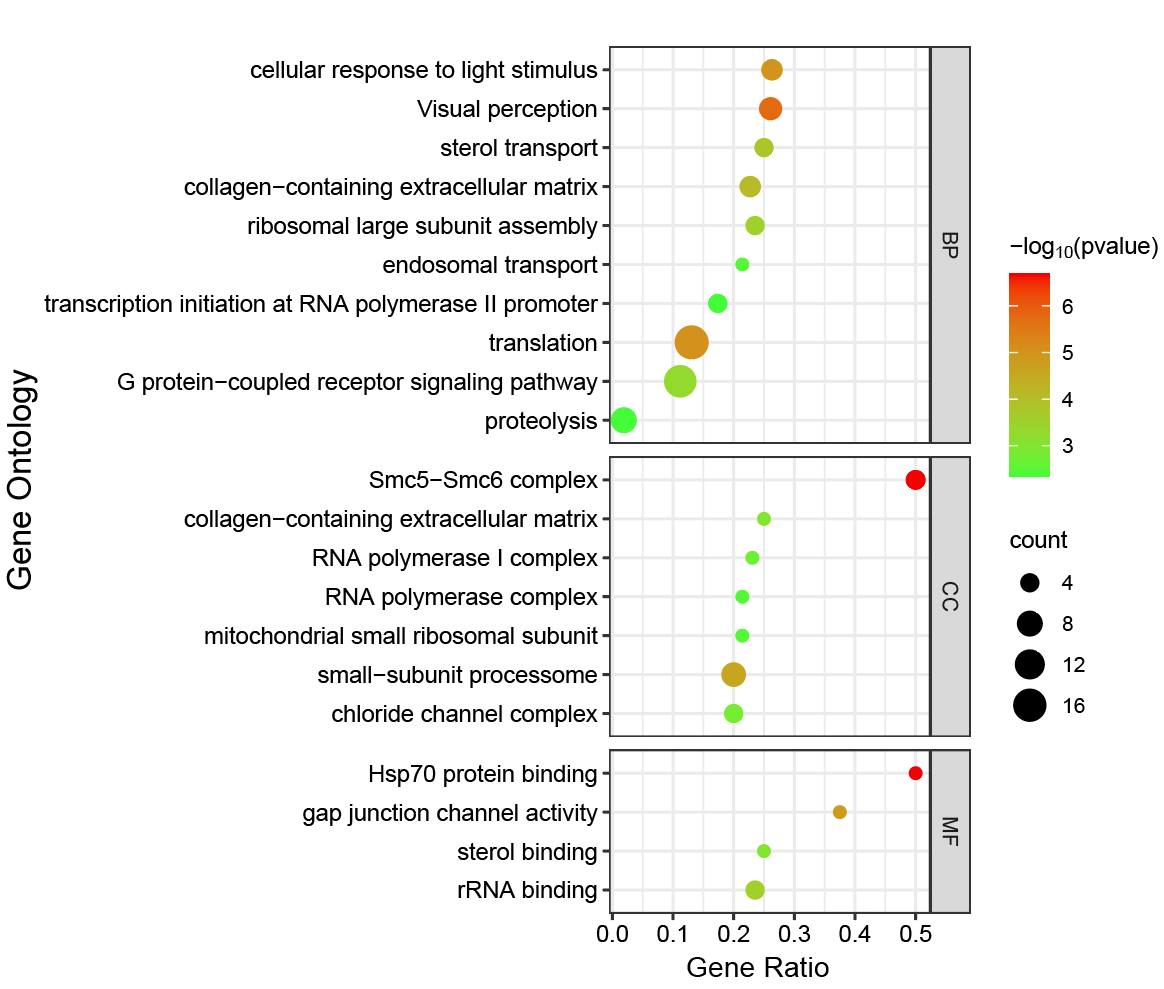
